# Fast Fourier Transform is a training-free, ultrafast, highly efficient, and fully interpretable approach for epigenomic data compression

**DOI:** 10.1101/2025.09.03.669431

**Authors:** Max Ward, Bac Dao, Amitava Datta, Zhaoyu Li

**Affiliations:** School of Physics, Mathematics, and Computer Sciences, University of Western Australia, Australia; School of Biomedical Sciences, University of Western Australia, Australia

## Abstract

Improving the efficiency of data compression remains essential for feature selection and data modelling. Current approaches for compressing epigenomic/genomic data highly rely on autoencoder that requires substantial computing resources, parameter fine-tuning, training, and time. Here, we developed a training-free, Fast Fourier Transform (FFT)-based method, for data compression with high efficiency and full interpretability. Our FFT method compresses epigenomic data of histone modification up to 1,000-fold while still maintaining high reconstruction fidelity (cosine similarity, 99.7%), does not require any training and completes ultrafast within 70 milliseconds on GPU or 20 seconds on CPU opposite to extensive training in hours/days for autoencoder on GPU/CPU, and offers full interpretability of compressed features from frequency components of original signals in contrast to the uninterpretable “black box” from autoencoder. This enables high accuracy in the classification model prediction (AUC, 0.960). Thus, our novel FFT method represents a major paradigm shift in data compression.

## Introduction

The Fourier Transform (FT) has long been established as a robust method for compressing and processing image data and sound signals^1-11^. The Fast Fourier Transform (FFT), first introduced by James W. Cooley and John W. Tukey, is an efficient algorithm for computing the Discrete Fourier Transform (DFT) and its inverse IDFT^12^. By factorizing the DFT matrix, the FFT significantly accelerated the computation and reduced the complexity to *O* (*n* log *n*)^12^, which is crucial for processing large datasets. FFT has been widely applied in various fields related to audio and image compression but never been used for processing epigenomic data.

The landscape of epigenetic research has been transformed by next-generation sequencing technologies to map genome-wide locations of epigenetic signals and regulators or epigenomic data, for instance, mapping genome-wide locations of histone modifications by chromatin immunoprecipitation coupled high-throughput sequencing (ChIP-Seq)^13^. The scale of these large epigenomic data is ideal for data modelling using machine learning. While these advances have provided unprecedented insights into chromatin states, massive datasets were also generated, posing significant computational and analytical challenges. This necessitates the application of effective data compression methods for handling these large datasets.

Many methods have been sought to project high-dimensional genomic and epigenomic datasets into lower-dimensional representations, such as the principal component analysis (PCA) integrated with sliced inverse regression^14^ or partial Cox regression^15^, an L0 segmentation-based method^16^. Recently, deep learning models, particularly autoencoders, have been heavily explored for epigenomic data compression, such as an Epi-LSTM-based autoencoder^17^, a 1D-convolutional variational autoencoder framework (ConvNet-VAEs)^18^, and a 300-fold compression approach but only with 60% accuracy^19^. Despite the potential, deep learning models like autoencoders require extensive computing resources of large computing units of GPUs, prolonged training time, careful hyperparameter tuning, and multiple times of tuning and training, resulting in significant challenge to be implemented in the large-scale studies. Autoencoders are widely considered to be a “black box” with limited interpretability. Recently, many efforts have been employed to increase the interpretation of autoencoder models with assisting from other approaches, such as removing non-linearity in the decoder^20^ and introducing dual embedding spaces with a weak correspondence term^21^. However, both approaches achieve only partial interpretability due to non-linear transformations in portions of their architectures and mathematical abstractions of their latent representations that require significant effort to connect to biological meaning. These limitations underscore the need for a more efficient, interpretable, and computationally tractable alternative.

Given that histone modification signals can be represented as continuous waveforms, we hypothesized that FFT could provide an efficient compression solution for epigenomic data. Here, we introduce a novel FFT-based method that achieves effective compression of histone modification data at single-nucleotide resolution, addressing the computing and interpretability limitations of current approaches.

## Results

### FFT transforms epigenomic data into fully interpretable frequency components

Briefly, we developed a PyTorch-based FFT approach for epigenomic data compression with the frequency annotation of each original signal (refer to details in Methods). The dataset used in this study was collected from genome-wide H3K4me3 signals of ChIP-Seq data in cell-free plasma samples from 88 healthy donors and 128 colorectal cancer patients from a previous report^22^.

To further expand the scale of data, we calculated the intensity of H3K4me3 signals at single-nucleotide resolution (details in Methods) across a genomic region of 300 kilobase (kb) surrounding the locus of a housekeeping gene, *GAPDH*, and then plotted the signal intensity versus relative genomic positions (Fig. 1a), where the peaks represent the enriched areas of H3K4me3 signals, indicating potential regulatory regions near active transcription start sites. These signals showed characteristic of H3K4me3 distribution patterns with both sharp, localized peaks and broader regions of enrichment (Fig. 1a). Next, we applied our FFT method to convert these H3K4me3 signals into the frequency data (details in Methods). Fig. 1b shows the frequency spectrum and magnitude of H3K4me3 signals with wide dynamic range obtained from the FFT analysis. The magnitude of frequency was calculated by the square root of the sum of the squared real and imaginary parts of the frequency component (details in Methods), representing the strength or contribution of each frequency component to individual original signals^23^. Along the frequency axis, low-frequency components (near 0) illustrate sharp, distinct peaks, representing genuine H3K4me4 signals in the genome, whereas high-frequency components (closer to 0.5) typically represent noise, rapid fluctuations, and background variations in the signals (Fig. 1b). More importantly, these compressed features from our FFT method directly correspond to those specific frequency components in the original signals, providing full interpretability of compressed data, which completely opposite to the uninterpretable “black box” representations from the autoencoder-based approaches.

**Fig. 1.**
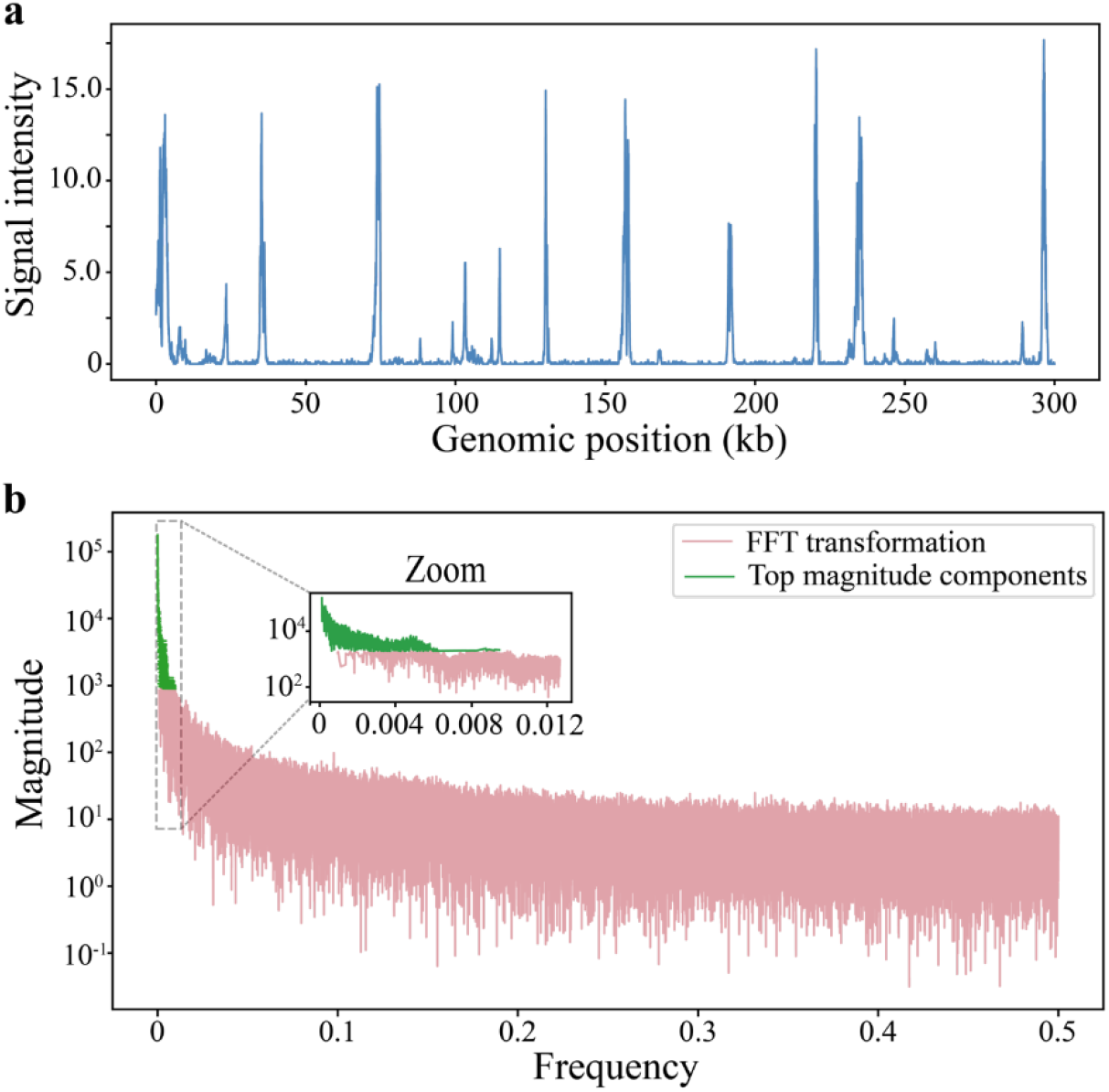
Transforming epigenomic signals of H3K4me3 into frequency data using FFT. **a**, The distributions of original H3K4me3 signal intensity from ChIP-Seq data across a 300 kb genomic region surrounding the GAPDH locus prior to the FFT analysis. kb, kilobase. **b**, The plot of magnitude and frequency obtained from the FFT analysis of H3K4me3 signals from (a). Zoom, a selected region highlighting 1,500 components with highest magnitude (green) for later signal reconstruction. The magnitude is displayed on a logarithmic scale.

### FFT achieves high efficiency of data compression

The high-magnitude components represent frequencies with the strongest influence on the signal structure. Using the same 300 kb genomic region in Fig. 1, we filtered the frequency spectrum to retain only the components with the highest magnitude (highlighted in green in Fig. 1b) while removing noise (details in Methods). Next, we used these high-magnitude components for data compression/reconstruction with our FFT method (details in Methods).

To evaluate our FFT method, we compared it with an autoencoder-based method reported previously^18^. Detailed autoencoder models architectures and the comparison pairs of selected features between FFT and autoencoder methods are described in Methods (Table 1 and 2 in Methods).

**Table 1.** Architectures oof autoencoder models.

| Compression ratio | Encoder Layers | Kernel Size (K) | Stride (S) | Decoder Layers |
| --- | --- | --- | --- | --- |
| 100X | Conv1D(1→16) | 6 | 2 | ConvTranspose1D(16→1) |
|  | Conv1D(16→32) | 6 | 5 | ConvTranspose1D(32→16) |
|  | Conv1D(32→64) | 6 | 5 | ConvTranspose1D(64→32) |
|  | Conv1D(64→1) | 6 | 5 | ConvTranspose1D(1→64) |
| 300X | Conv1D(1→16) | 6 | 5 | ConvTranspose1D(16→1) |
|  | Conv1D(16→32) | 6 | 5 | ConvTranspose1D(32→16) |
|  | Conv1D(32→64) | 6 | 4 | ConvTranspose1D(64→32) |
|  | Conv1D(64→1) | 5 | 3 | ConvTranspose1D(1→64) |
| 1000X | Conv1D(1→16) | 10 | 5 | ConvTranspose1D(16→1) |
|  | Conv1D(16→32) | 8 | 5 | ConvTranspose1D(32→16) |
|  | Conv1D(32→64) | 5 | 4 | ConvTranspose1D(64→32) |
|  | Conv1D(64→1) | 10 | 10 | ConvTranspose1D(1→64) |

**Table 2.**
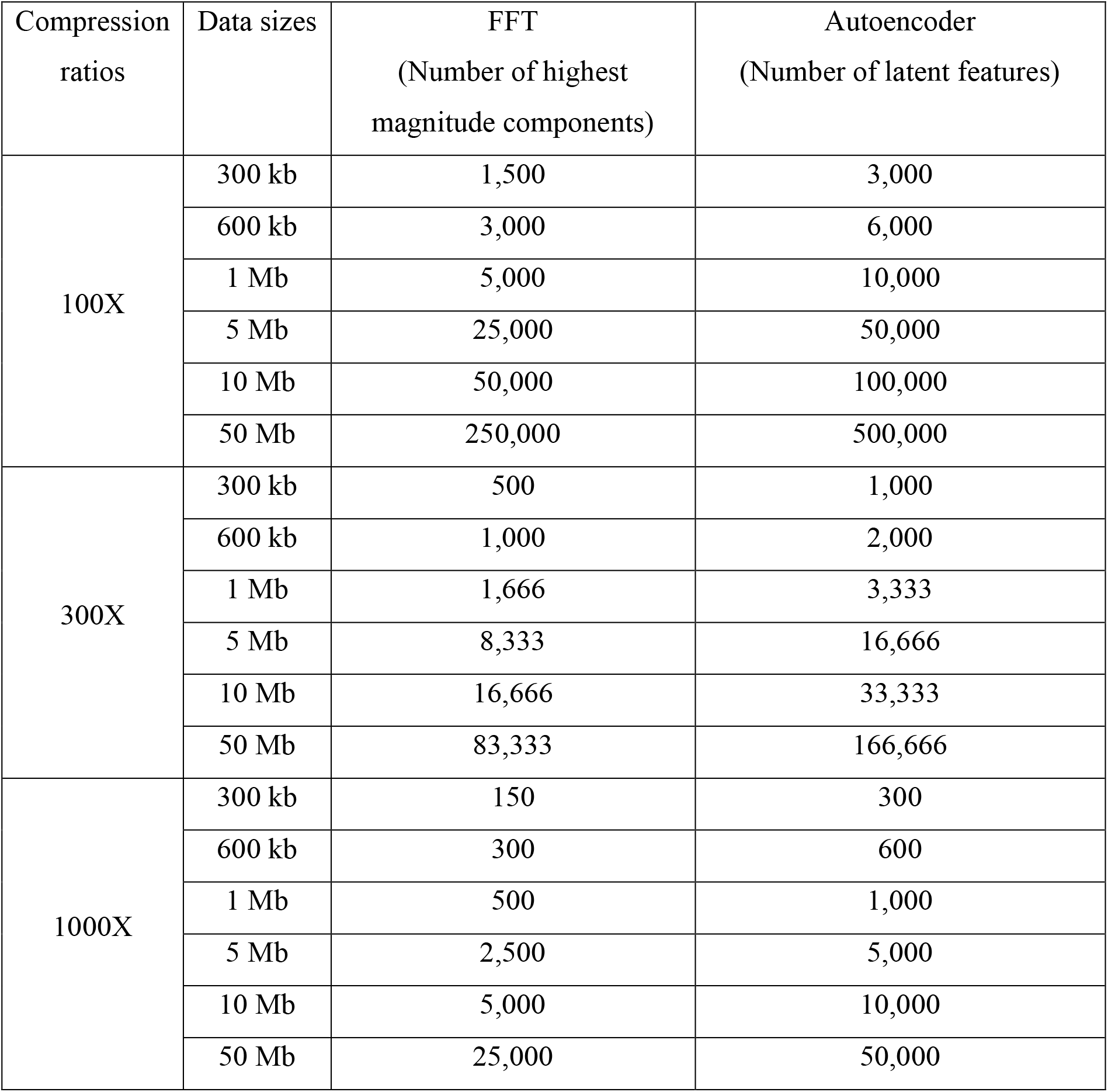
Features extraction parameter across multiple data sizes and compression ratios using FFT and autoencoder.

An example of reconstructed and original data of the same region of 300 kb from Fig. 1 at the 100-fold compression with the original signal intensity (blue) alongside two reconstructed signals, FFT (green) and autoencoder (orange) is shown in Extended Data Fig. 1a. We also extended our analysis to lager genomic regions from 300 kb to 50 million bases (Mb) (Extended Data Fig. 1,2) and further to individual chromosomes of the entire human genome (Fig. 2).

**Fig. 2.**
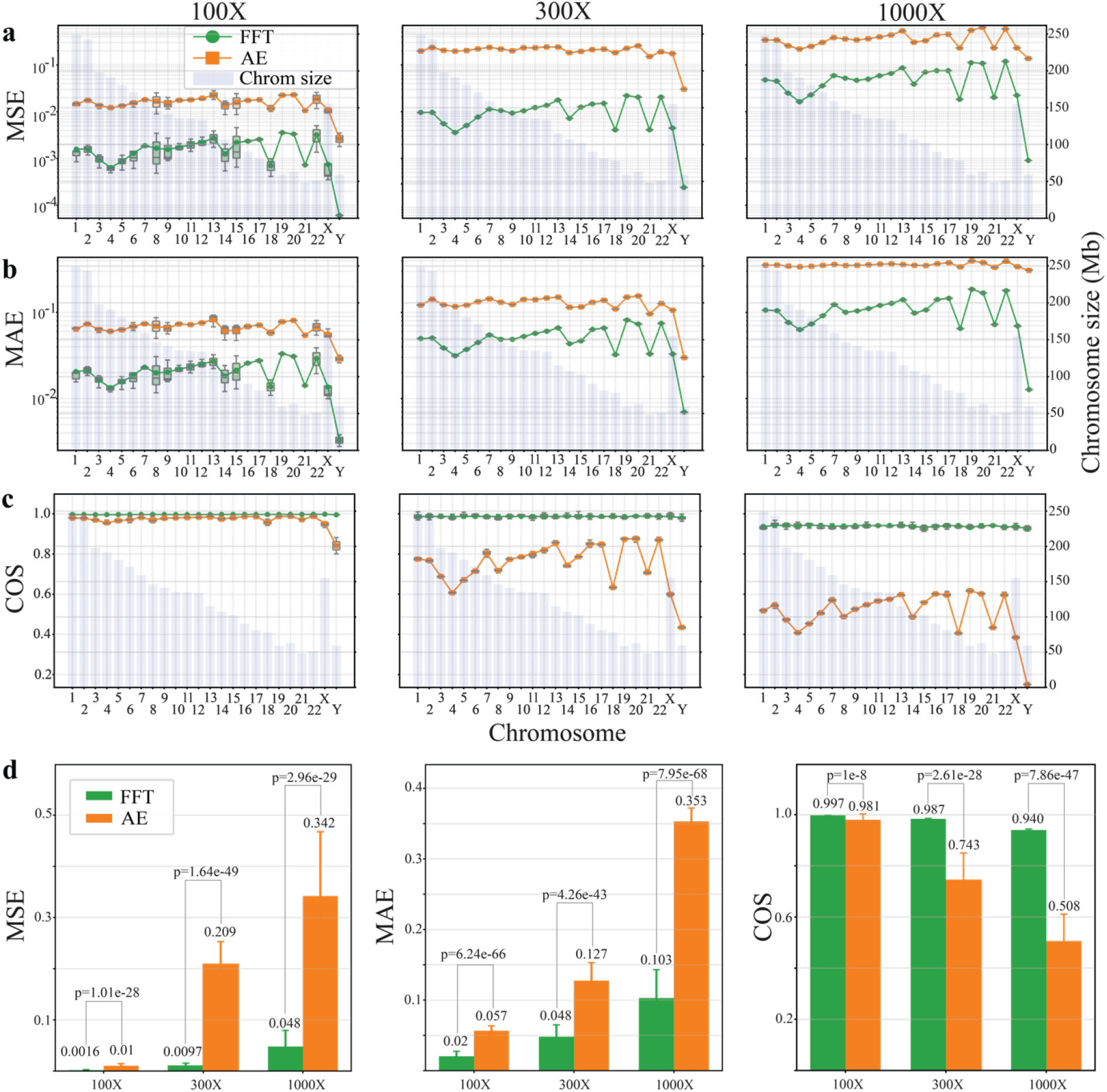
Chromosome-specific performance analysis using FFT and autoencoder methods to compress H3K4me signals across each chromosome with various compression ratios. **a,b,c**, Superior performance of our FFT method (green) to the autoencoder method (orange) in data compression of each consecutive 50 Mb segment of genomic regions across individual chromosomes as indicated by lower mean squared error (MSE), lower mean absolute error (MAE) and higher cosine similarity (COS) between reconstructed and original H3K4me3 signals with compression ratios of 100, 300, and 1000 folds (100X, 300X, and 1000X). Line plots demonstrate the mean value of performance metrics across individual chromosomes. Box plots illustrate the distributions of performance metrics across all 50 Mb genomic segments within each chromosome, with each box showing the interquartile range and median values. Grey bars represent chromosome sizes (right y-axis). FFT, Fast Fourier Transform; AE, autoencoder; Mb, million bases. **d**, Average of MSE, MAE, and COS across the entire genome. P values were calculated from comparison between FFT and AE metrics using *t*-test.

After data compression, three performance metrics, mean squared error, mean absolute error, and cosine similarity, between reconstructed and original data were calculated to evaluate the performance of both methods (details in Methods). The analysis of selected genomic regions from 0.3 to 50 Mb showed that consistently lower values of both mean squared error and mean absolute error from the FFT analyses compared to the autoencoder ones while the decrease in both values following the increase in the data sizes for both methods; the cosine similarity was constantly higher in our FFT analyses than the autoencoder (Extended Data Fig. 2), indicating higher fidelity of reconstructed data. Therefore, we used the 50 Mb as the segment size for genome-wide analysis.

At the genome-wide scale, we conducted the parallelled analysis of H3K4me3 signals at single-nucleotide resolution for all consecutive 50 Mb segments across each chromosome using both our FFT and the autoencoder methods with three compression ratios of 100, 300, and 1,000 folds (Fig. 2). Specifically, with the 100-fold compression, slight variations in the values of mean squared error and mean absolute error across all 50 Mb segments within the same chromosome were observed from both methods but diminished substantially at higher compression ratios (Fig. 2a,2b), indicating highly consistent results across all segments. Nevertheless, our FFT method still generated consistently and significantly lower values of mean squared error and mean absolute error in all chromosomes with all three compression ratios than those from the autoencoder method (Fig. 2a-2d). These results are aligned with our findings in smaller genomic regions (Extended Data Fig. 1,2), indicating that indicating the better signal reconstruction accuracy with minimal background noise of our FFT method compared to the autoencoder one.

The cosine similarity analyses also showed superior performance from our FFT method to the autoencoder one (Fig. 2c), exactly like the patterns observed from smaller genomic regions (Extended Data Fig. 2). More importantly, our FFT method showed consistent and super-higher cosine similarity (on average 0.997 for 100X, 0.987 for 300X, and 0.940 for 1,000X) in all chromosomes within each compression ratio, indicating the dependence on the compression ratios but independent of chromosome sizes; in contrast, the autoencoder method showed large variations of lower values of cosine similarity, particularly in those higher compression ratios of 300X and 1,000X (Fig. 2c,2d). These results indicate that our FFT method could capture nearly all features in the original data during data reconstruction. This high fidelity of data compression could be due to the full interpretability of compressed features from the frequency components (Fig. 1b).

### FFT does not require any training and runs ultrafast

Of a unique advantage was that our FFT method for data compression did not require any training. In contrast, autoencoder models usually require extensive time of hours to days to train the models depending on the data size and also multiple rounds of training. Fig. 3a defines the inference time and total time used for both FFT and autoencoder methods. The training time shown for autoencoder is only one-time training required initially but does not include potential extensive and multiple rounds of tuning and training afterwards. Total time represents the time required for completing the entire process of data reconstruction for each method. For FFT, the total time includes inference time plus reconstruction time. For autoencoder, the total time includes the time for training, inference time, and decoding phases together (Fig. 3a). Data related to inference time are shown in Extended Data Fig. 3.

**Fig. 3.**
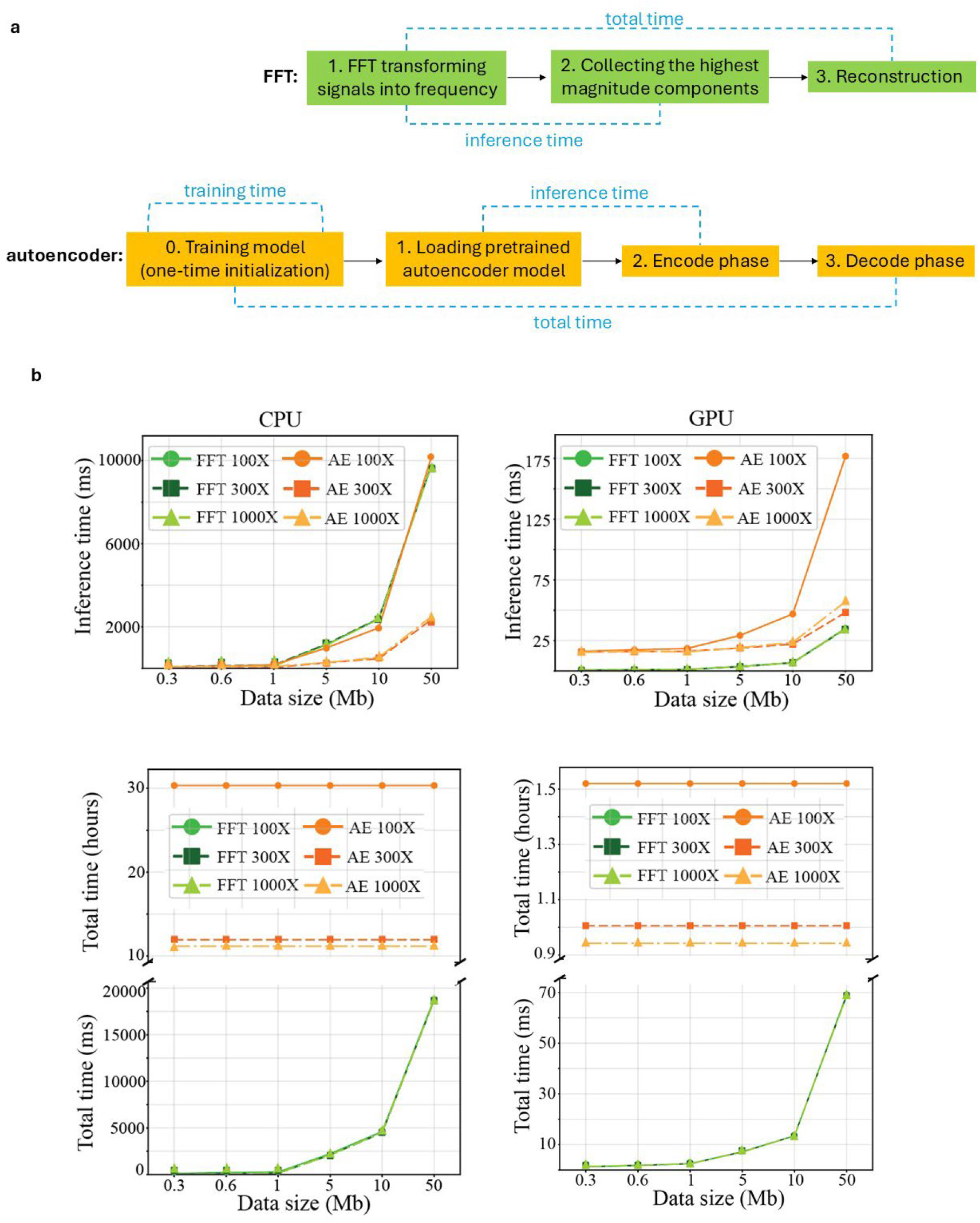
Comparison of the running time between FFT and autoencoder methods. **a**, Schematic workflow for our FFT and the autoencoder methods with definitions of inference time and total time. **b**, Performance comparison of both our FFT and the autoencoder methods running on CPU and GPU. The discontinued y-axis for total time includes milliseconds (ms) for FFT (bottom) and hours for autoencoder (top). 100X, 300X, and 1000X, 100, 300, and 1000 folds of compression; Mb, million bases.

We examined both our FFT and the autoencoder methods on both CPU and GPU (Methods). Shockingly, our FFT method was ultrafast and the FFT-based data compression was finished within 70 milliseconds (68.10 – 68.69 ms) across all different sizes of genomic regions from 0.3 to 50 Mb with all three compression ratios on GPU; in contrast, the autoencoder analysis required about 0.94-1.52 hours to complete with one round of training (Fig. 3b), representing at least 49,264 times of faster (0.94 hour/68.69 ms) of our FFT method than the autoencoder one on GPU. When running on CPU, our FFT compression was completed within 20 seconds (18.80 – 18.82 seconds) whereas the autoencoder analysis with one round of training required about 11 to 30 hours to complete (Fig. 3b). Thus, even though our FFT method runs on CPU, it is still at least 179 times (0.94 hour/18.82 seconds) faster than the autoencoder one running on GPU.

Notably, when analysing smaller genomic regions of less than 1 Mb, our FFT method basically completed at no time for all three compression ratios on both GPU and CPU and the total time were almost identical across all genomic sizes and three compression ratios on GPU and CPU, respectively (Fig. 3b). Such identical total time were also observed for the autoencoder method (Fig. 3b), given that the training time of hours accounts for the majority of the total time.

In summary, our FFT method dramatically outperformed the autoencoder one by delivering highly efficient data compression with substantially less computing time and computing resources.

### FFT retains key biological features in highly compressed data

To evaluate whether FFT-compressed data could be effectively utilized for the downstream analytical tasks, we developed a classification model for distinguishing between healthy and colorectal cancer samples using XGBoost^24^. To assess this model, we examined genomic regions associated with two key loci: *CDX1* (Caudal Type Homeobox 1) and *ELF3* (E74 Like ETS Transcription Factor 3). *CDX1* is an important tumour signature gene for colorectal cancer^25,26^. *ELF3* showed significant enrichment of H3K4me3 signals in colorectal cancer samples from this epigenomic dataset but was not previously identified as a tumour signature in the original study^22^.

We analysed H3K4me3 signals at these selected regions surrounding *CDX1* and *ELF3* using compressed data from both FFT and autoencoder methods. For the autoencoder analysis, we utilized latent feature values, while for the FFT method, we split complex numbers into two components and flattened them into a single one-dimensional array. Classification was performed using an XGBoost^24^ model with five-fold cross-validation. The dataset of total 216 samples (88 healthy donors and 128 colorectal cancer) was randomly split into training set (80%, n=172) and testing set (20%, n=44). We assessed the model performance using Receiver Operating Characteristic (ROC) curves, Area Under the Curve (AUC), and Average Precision (Extended Data Fig. 4a) with algorithms from scikit-learn^27^.

**Fig. 4.**
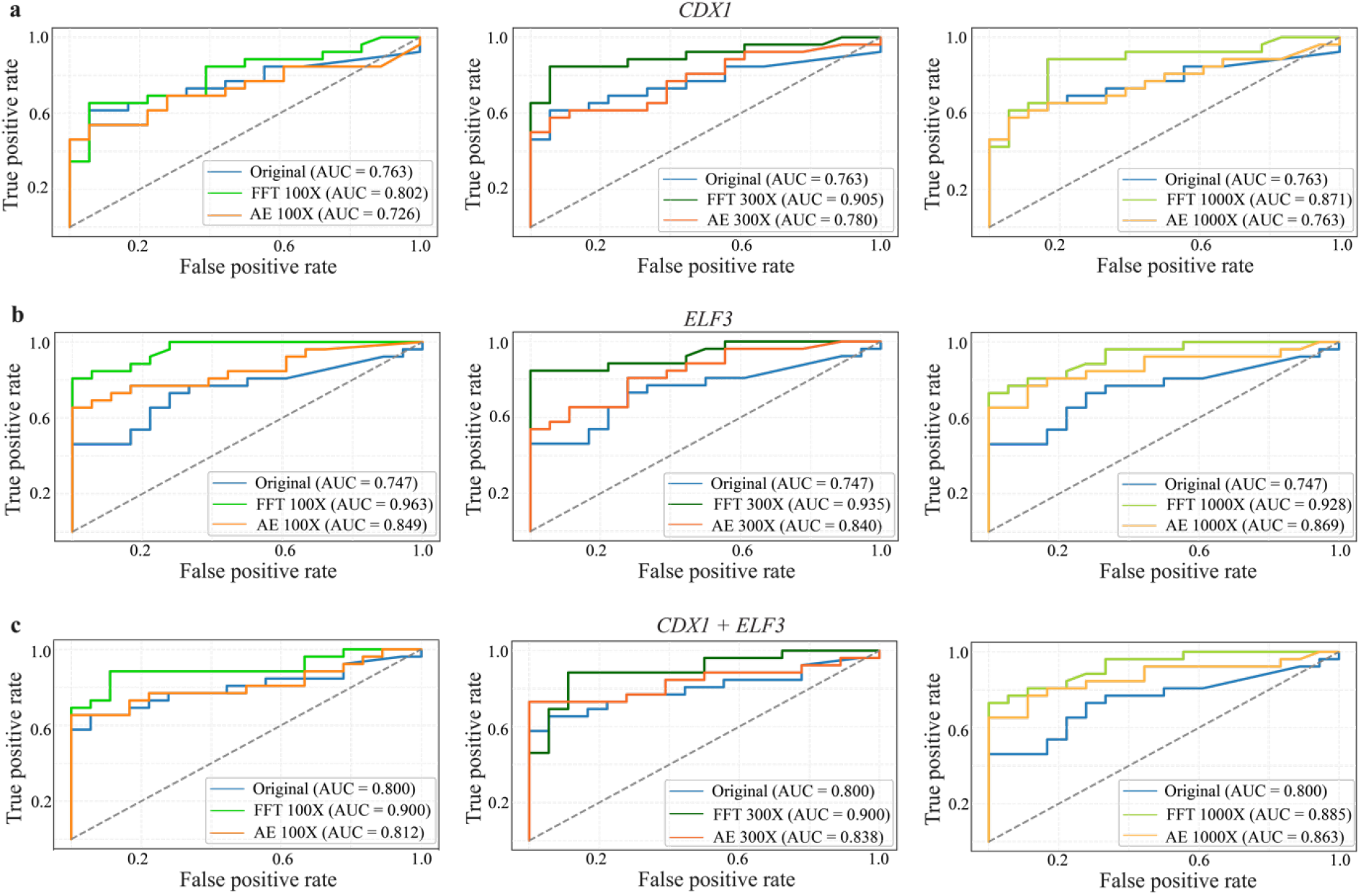
Comparison of the classification performance between original, FFT-compressed, and autoencoder-compressed H3K4me3 signals at genomic regions associated with *CDX1* and *ELF3*. **a,b,c**, Receiver-operating characteristic curves (ROC) for comparing the classification performance in distinguishing colorectal cancer samples from healthy samples using uncompressed original data (blue), FFT-compressed data a (green), and autoencoder-compressed data (orange) from genomic regions associated with *CDX1* (a), *ELF3* (b), and combined regions from both genes (c) with 100X, 300X and 1000X compression ratios.

The models built on the FFT-compressed data outperformed both models from the original uncompressed data and the autoencoder-compressed data at all three selected genomic regions across all compression ratios, as evidenced by higher AUC values (Fig. 4). For *CDX1*, the FFT-based models reached AUC values of 0.802, 0.905, and 0.871 were all greater than the AUC values of 0.726, 0.780, and 0.763, below the standard threshold of 0.8, from the autoencoder-based models for three compression ratios of 100X, 300X, and 1,000X, respectively (Fig. 4a). Therefore, the classification models built upon our novel FFT method showing much better performance further demonstrate that this epigenetic signature can serve as a robust biomarker for colorectal cancer.

More importantly, in our analysis, *ELF3* emerged as a novel and the most discriminative biomarker, achieving exceptionally higher AUC values of 0.963, 0.935, and 0.928 in the FFT-based models compared to 0.849, 0.840, and 0.869 from the autoencoder-based models after 100, 300, and 1,000 folds of compression, respectively (Fig. 4b). This finding supports previous studies indicating *ELF3* overexpression as a diagnostic marker ^28^ and its association with poor prognosis for colorectal cancer^29^.

The combination of both markers yielded robust classification performance from the FFT-based models (AUC = 0.90) (Fig. 4c). Similar results were also observed from the combined regions associated with top 54 genes (Extended Data Fig. 4b). This demonstrates that our FFT method is powerful not only for analysing the region associated with a single gene but also for integrating multiple regions spanning different chromosomes, paving the way for genome-wide classification.

## Discussions

In summary, we developed a novel, training-free, ultrafast, highly efficient, and fully interpretable FFT approach for data compression. The frequency data generated from our FFT method provides full interpretability with signal strength, position and periodicity. Therefore, such full interpretability allowed us to obtain accurate prediction of downstream targets from the classification modelling with high AUC values, shedding light on promising clinical applications. This approach could achieve up to a thousand fold of data compression while still preserving key biological features. Notably, compared to autoencoder-based approaches, our FFT method demonstrates superior accuracy and compression ratios, offering a more versatile, scalable, and fully interpretable alternative to neural network approaches for epigenomic data analysis. Moreover, our approach does not require any condition-specific training, enhancing its applicability across diverse datasets, particularly valuable in resource-constrained environments where GPU availability is limited. This innovation represents a major paradigm shift in data compression and also a significant advancement in the computational analysis of epigenomic data, enabling more comprehensive and nuanced investigations of chromatin states.

Our study is the first to apply FFT to compress epigenomic data, though until very recently, FFT started to be used for processing transcriptomic data^30,31^. Our FFT method can capture subtle differences in histone modification data at single-nucleotide resolution superior to the traditional ‘binning’ strategy for the epigenomic analysis^32,33^. More importantly, our FFT approach holds promise for future work exploring the application of this method to other types of genomic and epigenomic data, potentially revealing frequency domain characteristics of biological significance across different regulatory landscapes.

## Data and code availability (code is deposited at personal GitHub account)

The epigenomic data were collected from the Zenodo repository: https://doi.org/10.5281/zenodo.4277001 as published by Sadeh et al^22^.

The codes are deposited at GitHub temporarily and will be moved to a permanent repository after acceptance. GitHub: https://gitfront.io/r/bacdao/XzG83cqHu67h/FFT/

## Acknowledgement

This study was partially supported by the Research Project grant from Cancer Council WA to Z.L., A.D., and M.W.

B.D. was supported by an Australian Government Research Training Program (RTP) Scholarship from the University of Western Australia.

## Author contributions

Project conceptualization, methodology, supervision, writing, funding acquisition: Z.L., M.W., A.D.; investigation: Z.L., M.W., A.D., B.D.

## Methods

### Data processing

We obtained genome-wide histone modification data of H3K4me3 from the Zenodo repository as published by Sadeh et al^1^. This dataset was generated using cfChIP-Seq technology to profile histone modifications in cell-free plasma samples from 88 healthy donors and 128 colorectal cancer (CRC) patients.

For the single-nucleotide resolution processing, we defined a genomic sequence *G* of length *n* base pairs, where each position *g ∈ G* is indexed from 1 to *n*. The set of H3K4me3 peaks *P* consists of tuples *(s,e,v)*, where s and e denote the start and end positions, respectively, and *v* represents the peak intensity value. The signal intensity *S(g)* at genomic position *g* is defined as:

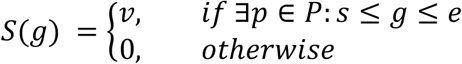

This formulation ensures that genomic positions within peak regions inherit their corresponding peak intensity values, while positions outside these regions are assigned zero intensity.

### FFT Implementation

We developed a PyTorch-based Fast Fourier Transform (FFT)^2^ approach for compressing epigenomic signals of histone modification. Given a signal vector *x[n]* of length *L*, the FFT transformation *F(k)* is computed as:

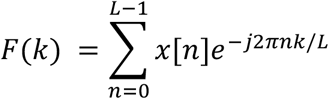

for *k ∈ [0, L/2 + 1]* in the real FFT case.

The magnitude of each frequency component was then computed using^24^:

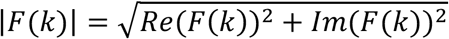

For a given compression ratio *r*, we retained the *m=L/(2r)* frequency components with the highest magnitudes, achieving a compression ratio of *r/2* by storing *2m* float values (corresponding to *m* complex numbers).

For signal reconstruction, a frequency domain array *F′(k)* of length *L/2+1* was constructed, populated with the stored highest-magnitude components, and subjected to inverse FFT:

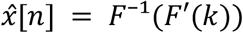

This approach was applied across data sizes *L ∈* {300k, 600k, 1M, 5M, 10M, 50M} base pairs (bp) with r *∈* {100, 300, 1000}.

The frequency spectrum spans from 0 to 0.5 in normalized frequency units, a direct consequence of the Nyquist-Shannon sampling theorem^3,4^. This theorem establishes that a discretely sampled signal can only represent frequency components up to half the sampling rate without aliasing. Since our data is sampled at single-nucleotide resolution (sampling rate = 1), the highest representable frequency is precisely 0.5 cycles per nucleotide (Fig. 1b).

### Comparative analysis with autoencoder

To benchmark our FFT-based method, we implemented an autoencoder model with identical compression ratios (100X, 300X, and 1000X). For model training, we used a mini-batch size of 64, Adam optimizer (learning rate=0.0001) with ReduceLROnPlateau scheduler (factor=0.1, patience=10). The autoencoder architecture incorporated 4 convolutional layers in both encoder (1→16→32→64→encoding_dim) and decoder (encoding_dim→64→32→16→1) paths. The architecture utilized Batch Normalization after each convolutional operation and ReLU activation functions in all layers except the final output. The model was trained for a maximum of 1000 epochs across 100 distinct genomic regions, each spanning 300 kilobase pairs to capture diverse histone modification signals. Early stopping (patience=50) was implemented to prevent overfitting. Training was performed across 2 NVIDIA Tesla V100-PCIe-32GB GPUs using PyTorch’s DataParallel for acceleration, with data split into training and validation sets (80%/20%). For each compression ratio, the kernel sizes and strides were adjusted accordingly, and the model with the lowest validation loss was preserved for subsequent analyses.

The detailed architecture of these models, including the number of convolutional layers, kernel sizes, stride values, and channel dimensions, is provided in Table 1. Model performance was subsequently evaluated on independent genomic regions excluded from the training phase.

A comparative assessment was conducted using three quantitative metrics: mean squared error (MSE), mean absolute error (MAE), and cosine similarity (COS), measuring the fidelity of reconstructed signals relative to the original data. These evaluations were performed across multiple data sizes (300 kb, 600 kb, 1 Mb, 5 Mb, 10 Mb, and 50 Mb) to assess the scalability and robustness of each method.

MSE and MAE were used to quantify the overall pointwise difference between the original and reconstructed data. Lower values of MSE and MAE indicate better reconstruction accuracy across the entire dataset. However, these metrics do not capture the similarity in patterns or trends between the original and reconstructed data. In some cases, the reconstructed data may have small MSE and MAE values, suggesting good pointwise accuracy, but still exhibit different patterns or trends compared to the original data. To address this limitation, cosine similarity was employed to evaluate the similarity between the original and reconstructed data vectors in terms of their orientation, regardless of their magnitudes. Higher cosine similarity values (closer to 1) indicate a higher degree of pattern preservation and suggest that the reconstructed data closely follows the trends and structure of the original data.

The MSE was computed as:

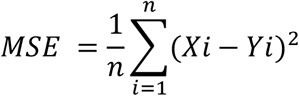

The MAE was computed as:

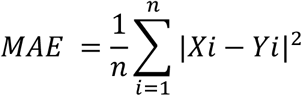

where Xi represents the reconstructed signal and Yi represents the original signal. COS was computed as:

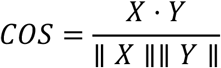

where *X* · *Y* is the dot product, and ∥ *X* ∥, ∥ *Y* ∥ are the Euclidean norms of the respective signals.

Since the Fourier transformation of real-valued signal intensities produces complex numbers (requiring twice the memory storage), after selecting the numbers of highest-magnitude frequency components for the FFT analysis, we used the double numbers of latent features for the autoencoder analysis on the same genomic region (Table 2). This approach ensures a fair comparison by maintaining consistent memory requirements across different compression levels.

For example, at the 100-fold compression, we configured the autoencoder with hidden layers to produce 3,000 latent features (the original 300,000 dimensions divided by 100 folds of compression ratio); however, for our FFT method, we retained the top 1,500 highest-magnitude components (highlighted in green in Fig. 1b) to ensure equivalent memory storage requirements, as each complex FFT component requires twice the memory of a real-valued autoencoder feature. With the 100-fold compression, this experimental design allowed us to directly compare the reconstruction accuracy between the 3,000 autoencoder latent features and the 1,500 FFT highest-magnitude components (highlighted in green in Fig. 1b) for the selected 300 kb region. We also applied this to all other selected genomic regions with other paired comparison features between our FFT and the autoencoder methods for different sizes of genomic regions with three compression ratios of 100-fold, 300-fold, and 1000-fold (Table 2) to evaluate the optimal balance between storage reduction and information retention.

kb, kilobase; Mb, million bases.

### Chromosome-wise comparative analysis

To rigorously evaluate the consistency of both FFT and autoencoder methods across variable genomic contexts, we partitioned each chromosome into contiguous non-overlapping 50 Mb segments and analysed them at single-nucleotide resolution. For each segment, we quantified reconstruction fidelity using MSE, MAE, and COS.

### Classification Model

To evaluate the downstream utility of our compression methods, we developed a classification framework to distinguish between healthy and colorectal cancer samples using three datasets: (i) original signals of H3K4me3 from ChIP-Seq data, (ii) FFT-compressed signals, and (iii) autoencoder-compressed signals. Our analysis was focused on those genes with differential signals of H3K4me3 between healthy and cancer samples. For each gene, we analysed the entire gene body plus an additional 2 kb upstream of its transcription start site at single-nucleotide resolution to capture the complete signature of H3K4me3 signals.

For classification, we employed XGBoost^5^ with GPU acceleration, configured with 500 estimators, a maximum tree depth of 50, and a learning rate of 0.1 with five-fold cross-validation. The dataset was stratified into 80% training and 20% testing sets. Model performance was evaluated using receiver operating characteristic (ROC) curves, area under the curve (AUC) scores, and average precision (AP) using scikit-learn^6^.

### Implementation

All computational analyses were performed using PyTorch 2.5.1 and Python 3.11.10. Both the autoencoder architecture (implemented with PyTorch’s neural network modules) and FFT operations (utilizing PyTorch’s native implementation) were accelerated on NVIDIA Tesla V100-PCIe-32GB GPUs at Kaya supercomputing cluster owned by the University of Western Australia. For testing time performance between FFT and autoencoder on CPU, evaluations were performed on a single core CPU (Intel(R) Xeon(R) Gold 6254 CPU @ 3.10GHz). The classification model was implemented using XGBoost 2.0.3 and executed on the same GPU architecture. Performance metrics including MSE, MAE, COS, ROC-AUC, and AP were calculated using scikit-learn 1.5.2^6^.

